# Functional Continuum of GABAergic Synaptic Dynamics Reflects Genetic Identities

**DOI:** 10.64898/2026.06.09.731181

**Authors:** Jade Poirier, John Beninger, Katalin Tóth, Richard Naud

## Abstract

The language of the brain is articulated by temporal patterns of neuronal activity, which individual synapses interpret through distinct forms of synaptic dynamics. While glutamatergic synapses have been proposed to use a handful of functionally distinct types of short-term plasticity (STP) that loosely align with genetic identities, the functional organization of GABAergic synaptic dynamics remains ambiguous. Here, we ask whether the diversity of GABAergic dynamics also clusters into discrete functional types or instead forms a continuum, and whether this functional organization also aligns with genetic identities. Using a combination of machine learning and synaptic modeling on the Allen Institute Synaptic Physiology Dataset, we present evidence that inhibitory dynamics organize into a functional continuum with overlapping modes corresponding to different genetic identities. This continuum spans Parvalbumin (Pvalb) to Vasoactive Intestinal Peptide (VIP) presynaptic neurons, with Somatostatin (Sst) neurons forming an intermediate class. Across the three classes, most inhibitory synapses showed either strong depression or a biphasic form of plasticity that is facilitating at higher stimulation frequencies and depressing at lower ones. However, a higher proportion of Sst and VIP synapses, compared to Pvalb synapses, exhibited either depression restricted to high frequencies or consistent facilitation. When paired with functional subtypes of excitatory synaptic dynamics, this inhibitory continuum completes a unified framework for understanding information flow in the cortical microcircuit, advancing long-standing efforts to explain circuit function by investigating the organization of its components.

## Introduction

Transient changes of individual synapses are fundamental to the brain’s ability to dynamically regulate information flow. Known as short-term plasticity (STP), these activity-dependent changes in synaptic strength typically persist for milliseconds to seconds, a timescale that aligns with many cognitive operations.^1-4^ While systems neuroscience has largely acknowledged the need to characterize neurons both in terms of their connectivity profiles and cellular subtypes to understand their functions within circuits,^5^ in-depth exploration of the distribution of short-term dynamics, particularly their alignment with cell types, remains comparatively incomplete. Because STP can transiently alter a synapse’s effective connectivity, such exploration is essential for advancing our understanding of circuit-level neural computation.

At the mechanistic level, STP has long been associated with presynaptic factors that transiently alter vesicle availability and release probability, giving rise to facilitation at synapses with initially low release probability and depression at those with high release probability.^4^ One such mechanism involves the accumulation of residual calcium in the presynaptic terminal,^6^ increasing vesicle release probability temporarily through the modulation of Ca^2+^-binding proteins.^7-11^ In contrast, depletion of the readily releasable pool during sustained activity is a well-established driver of depression.^4^ Retrograde messengers, such as endocannabinoids^12,13^, can also act on presynaptic receptors, biasing the synapse towards facilitation or depression. Additional postsynaptic factors have been described over the years: AMPA receptor desensitization^14^ and diffusion-trapping,^15^ as well as gradients of short-term dynamics along dendrites^16,17^ are examples of such postsynaptic contributions. Factors external to the synapse further expand this landscape; for instance, astrocytic clearance of glutamate can promote depression,^18^ while messengers released from surrounding synapses can modulate presynaptic release.^4^ While not exhaustive, this list of interacting biological mechanisms highlights how mechanistic variety naturally produces a broad repertoire of temporal responses that extend beyond simple facilitation or depression.

Echoing this diversity, numerous variants of STP have been described, such as biphasic STP,^19,20^ delayed burst-dependent STP,^21^ and sub- and supralinear facilitation.^22^ Short-term dynamics are also known to undergo long-term changes as part of normal developmental processes^23^ and in an activity-dependent manner,^24^ which has led to the hypothesis that STP may be governed by learning rules independent of those underlying long-term plasticity.^25^ These processes likely contribute to the considerable heterogeneity of short-term dynamics but also suggest the existence of a functional organization in which specific STP profiles are matched to circuit-level computational demands.

On one hand, it was proposed that STP may have filtering properties: short-term facilitation (STF) could serve as a high-pass filter, whereas short-term depression (STD) could serve as a low-pass filter.^26^ By responding preferentially to certain temporal patterns, synapses could also extract distinct signals from a shared input based on their STP profile, thereby decoding a multiplexed code.^27,28^ Relatedly, previous studies suggest that the diversity in short-term dynamics could form a reservoir of temporal filters,^29,30^ placing STP as a key player in learning temporal associations and contributor to the brain’s ability to parse and predict unfolding events.^31-34^ Despite these hypotheses and the prevalence of STP across the brain,^35,36^ there is still no consensus on its computational role. As it is, a major challenge in deciphering the function of STP lies in its heterogeneity, which exists both within and across genetically defined cell types.^37, 38^

Previously, Campagnola et al. used UMAP dimensionality reduction^39^ to examine this synaptic heterogeneity, finding two broad, well-separated groups: one excitatory and one inhibitory, with continuous variation of dynamics within each.^38^ However, because UMAP is a nonlinear embedding that preserves local neighborhoods but distorts global structures, it cannot determine whether discrete subtypes of excitatory or inhibitory dynamics exist, which requires explicit clustering and statistical tests. Later, Beninger et al. used a combination of synaptic modelling and machine learning to demonstrate that excitatory synapses from the same dataset organize into functional subtypes, which partly overlap with genetic cell identities.^40^ Such lines of investigation are vital for providing a framework to understand how different cells and circuits contribute to overall brain function.^5^ However, this framework remains incomplete without a detailed account of the functional organization of inhibitory synaptic dynamics in the cortical microcircuitry.

To address this gap, we analyzed the largest available paired-patch clamp synaptic dynamics dataset (i.e., the Allen Institute Synaptic Physiology Dataset).^38^ By fitting a flexible synaptic model, we obtained low-dimensional representations of inhibitory STP. Supervised prediction of presynaptic cell class from model parameters revealed above-baseline accuracy, indicating that inhibitory short-term dynamics reflect meaningful genetic information. At the same time, unsupervised methods did not yield discrete clusters beyond what would be expected from noise of similar variance structure to the data, suggesting that inhibitory STP does not segregate into defined categories. This contrasts with Beninger et al. previous results on excitatory synapses, which we found to aggregate into distinct subtypes while using a similar methodology as described here.^40^ Instead, both the fitted model parameters and the supervised prediction weights for the inhibitory dataset revealed a smooth, overlapping continuum spanning Parvalbumin (Pvalb) to Vasoactive Intestinal Peptide (VIP) presynaptic neurons, with Somatostatin (Sst) neurons forming an intermediate class. This organization provides a new framework for understanding how inhibitory synapses contribute to circuit-level computations through graded modes of short-term dynamics. Further, by completing an account of the functional organization of synaptic dynamics in the cortical microcircuit, this framework pairs with existing literature to culminate in a basic dictionary of the language of the brain.

## Results

Throughout this study, we used paired-patch recordings of GABAergic synapses obtained from the Allen Institute Synaptic Physiology Dataset, the largest dataset publicly available of its kind. In this dataset, up to eight neurons from mouse or human slices were selected per experiment for simultaneous whole-cell patch-clamp recording in current-clamp conditions. Each patched neuron was administered 12 pulses at fixed frequencies (10 to 200 Hz), with a delay period between the 8^th^ and 9^th^ pulses (Figure 1A). In mice, both pre- and postsynaptic cells were genetically identified for their transgenically defined subclass.^38,41-43^ After quality control steps (see Methods), 940 rodent and 69 human synapses remained, thus fulfilling the size requirements for proper training and testing of the machine learning algorithms detailed below.

**Figure 1.**
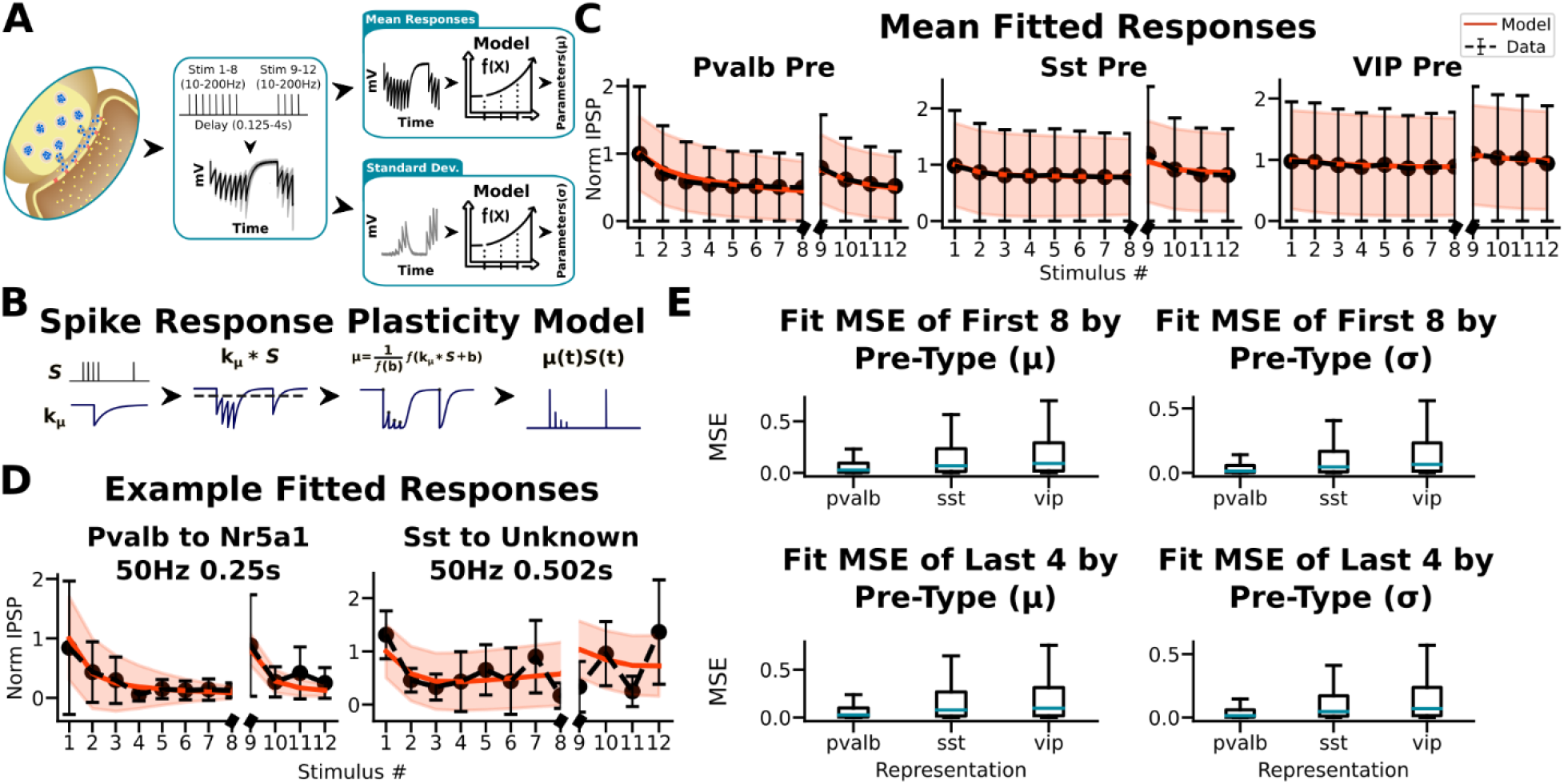
Spike Response Plasticity (SRP) model and fit quality on rodent data. **A**. Schematic of model fitting procedure, as described in Methods. **B**. Schematic of the SRP model. First, an input spike train is convolved with an efficacy kernel composed of exponentially decaying functions; then, a baseline parameter is added to the output, and the result is passed through a nonlinear (sigmoidal) function. The resulting signal sampled at the spike time makes the predicted efficacies. **C**. Mean SRP model fits (red) and data (black) for all synapses sharing the same presynaptic cell type (574 from Pvalb, 297 from Sst, and 69 from VIP). The error bars and shaded areas show the responses’ actual and predicted standard deviations, respectively. **D**. Example fits for two synapse pairs (a Pvalb→Nr5a1 synapse and an Sst→Unknown synapse) using specific stimulation protocols, corresponding to 50 Hz trains with 8th–9th inter-pulse delays of 0.25 s and 0.502 s, respectively. **E**. Mean squared error (MSE) of fit on the responses (left) and their standard deviation (right) to the first 8 stimuli before the delay and the last 4 stimuli after the delay, grouped by the presynaptic cell subclass. The teal-colored bars represent the median of each distribution.

### The Spike Response Plasticity (SRP) model captures diverse GABAergic STP

To investigate the functional organization of inhibitory dynamics through clustering, we first characterized dynamics at each synapse. Specifically, we fitted a flexible synaptic model (i.e., the Spike Response Plasticity (SRP) model)^44,45^ to recorded responses at each synapse, resulting in a condensed, low-dimensional representation of their dynamics in the form of model parameters. These parameters define the shape and baseline of either an efficacy (μ) or variance (σ) kernel, which is then convolved with an input spike train (i.e., the 12 pulses) (Figure 1A & 1B). Here, the kernels respectively capture the mean short-term dynamics, as well as the change in standard deviation of responses across pulses at a given synapse, with each kernel characterized by its own set of parameters. Once fitted to recorded responses, the model predicts how a synapse would respond to a given input spike train, according to its STP properties.

We applied standard optimization techniques^45^ to minimize the mean squared error (MSE) between predicted and recorded normalized IPSP amplitudes, and the MSE between predicted and recorded standard deviations of those amplitudes. Figure 1C showcases strong average fitting performance when pooling all pairs that share a presynaptic cell type (i.e., Pvalb, Sst, or VIP). The reasoning behind pooling across presynaptic types stems from evidence that, in inhibitory synapses, presynaptic types tend to be more predictive of their short-term synaptic than postsynaptic ones.^38^ Example fits in Figure 1D illustrates the wide flexibility of the model, capable of capturing both depression and a decrease in the SDs of responses over time (left) as well as a mixture of depression and facilitation on different timescales and an increase in the SDs across pulses (right). This fitting performance can be translated into mean MSEs, which are low across all synapses (μ: mean MSE and SEM of 0.1554±0.0021 and 0.1725±0.0036 over the first 8 stimuli and the last 4 stimuli; σ: mean MSE and SEM of 0.1108±0.0014 and 0.1157±0.0021 over the first 8 stimuli and the last 4 stimuli) (Figure 1E).

The strong average fitting performance and apparent flexibility of the model supports its use for the subsequent analyses.

### Inhibitory dynamics organize into a functional continuum

To determine whether subtypes of synaptic dynamics can be extracted from the dataset, we applied the OPTICS density-based clustering algorithm^46^ to the efficacy kernel parameters exclusively. Because the dataset is class-imbalanced, we performed bootstrap subsampling before clustering with equal numbers of synapses per presynaptic or postsynaptic subclass. Rodent and human synapses were clustered separately across a range of minimum cluster sizes, and for each bootstrap iteration we selected the clustering with the highest Silhouette Coefficient (SC), a metric of clustering quality (Figure 2A, upper branch).

**Figure 2.**
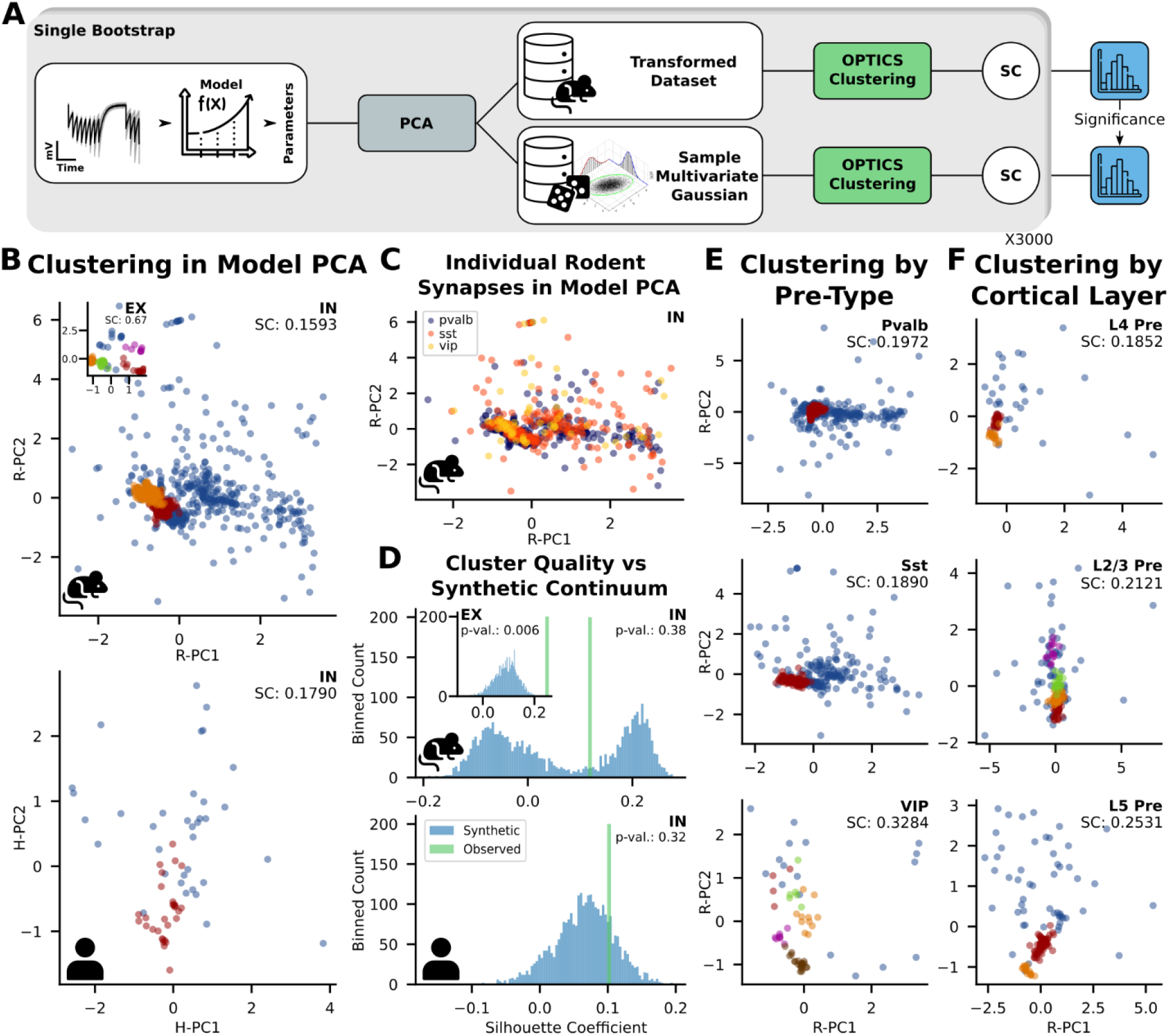
Inhibitory synaptic dynamics show no significant clusters in mouse and in human. **A**. Schematic demonstrating the clustering procedure onto the biological data (upper branch) as well as the one used to generate a noise distribution sharing a similar variance structure to the first. This process was repeated 3000 times, each iteration yielding a single Silhouette Coefficient. **B**. Rodent (top) and human (bottom) clustering, individual synapses being projected onto their respective PC1-PC2 spaces (i.e., R-PC=rodent principal component and H-PC=human principal component). Upper left corner: rodent excitatory dynamics groupings (PC1-PC2), showing distinct clusters. **C**. PCA projections of SRP parameters for inhibitory synapses colored by presynaptic genetic identities. **D**. Histogram of Silhouette Coefficients (a metric of clustering quality) computed by clustering class-balanced subsets of SRP parameters (“Mean Observed”: green) and noise distributions (“Synthetic”: blue). Minimum cluster sizes were selected on the basis that exactly two clusters would be generated. Rodent: p=0.38; Human: p=0.32. **E**. Clustering performed on SRP parameters, separated by presynaptic class **F**. Clustering performed on SRP parameters, separated by the presynaptic neuron’s laminar location. “SC”: Silhouette Coefficient.

Across all rodent inhibitory synapses, OPTICS identified two clusters with a low SC (SC=0.16). In human inhibitory synapses, it identified a single cluster with similarly low quality (SC=0.18) (Figure 2B). These values contrast sharply with those obtained for excitatory synapses in the same dataset (SC=0.67 for the five excitatory rodent clusters; SC=0.71 for the six excitatory human clusters).^40^ Figure 2C shows the overlapping dynamics between synapses with varying presynaptic cell types (i.e., Pvalb, Sst, or VIP). Moreover, inhibitory synapses showed extensive overlap across presynaptic cell types (Pvalb, Sst, VIP), with no clear subtype separation (Figure 2C). That said, a clustering and its associated SC can be influenced by a multitude of factors, such as the size of the dataset, its dimensionality and noisiness. Therefore, the significance of the clusters found in Figure 2B can’t be evaluated purely based on SCs.

Because OPTICS identifies density-based clusters, it will always predict the presence of at least two groupings (e.g., a dense region interpreted as a cluster and a set of noise points) regardless of the dataset’s underlying structure. Therefore, to assess the significance of the observed clusters, we compared measures of their quality to measures of quality from clusters inferred from synthetic data generated from a functional continuum (i.e., a model without distinct subtypes). To this end, we generated synthetic synapses by sampling from a multivariate Gaussian distribution whose variance structure matched the variance structure of the models fit to data from the real synapses, as captured by principal component analysis (Figure 2A, lower branch). We selected the first principal component as the primary axis of elongation for the multivariate Gaussian, then redistributed the remaining variance equally across the other axes so that the resulting distribution is more strongly stretched along one dimension while maintaining equal variance in the remaining directions. We could then compute SCs on clusters obtained from OPTICS applied separately to both sampled biological and synthetic data across a range of minimum cluster sizes.

For comparability between the biological data and its corresponding synthetic data, we selected only the highest SC from clustering results that yielded exactly two clusters at each bootstrap iteration. Doing so allowed us to avoid the problem of comparing SCs produced by drastically different clustering outcomes (e.g., cases where two clusters were identified versus cases with dozens of clusters), differences that arose from both our protocol and the unstable regime in which inhibitory-synapse clustering appeared to operate. Figure 2D shows the distributions of these SCs from the inhibitory rodent and human datasets, where in both cases the clustering quality did not show a statistical difference between the biological and synthetic data (p=0.38 for the rodent dataset; p=0.32 for the human dataset). In contrast, the quality of clusters of the excitatory rodent synapses was significantly better than the quality of clusters of their corresponding synthetic data (i.e., p=0.006). This is in accordance with previous results from Beninger et al. on the same synapses.^40^ As such, while findings from excitatory synapses support the notion that their dynamics separate into discrete subtypes, those related to inhibitory synapses are consistent with what would be expected from dynamics organizing along a continuum.

Since applying OPTICS globally to each dataset may overlook biologically relevant subtypes tied to specific cell types or cortical layers, we repeated the clustering process on targeted subsets of the rodent dataset. Clustering on subsets of rodent synapses sharing either a presynaptic cell type (Figure 2E) or presynaptic layer (Figure 2F) yielded relatively low SCs (SC_pvalb-pre_=0.20, SC_sst-pre_=0.19, SC_vip-pre_=0.33, SC_L4-pre_=0.19, SC_L2/3-pre_=0.21, SC_L5-pre_=0.25). In other words, while considering biologically relevant subsets of the dataset, we couldn’t identify meaningful subtypes of inhibitory dynamics.

### The dynamics of inhibitory connections align with presynaptic genetic identities

To assess whether features of synaptic dynamics can predict the transgenic class of the presynaptic (Pvalb, Sst, VIP) or postsynaptic (Nr5a1, Tlx3, Sst, Ntsr1, VIP, Sim1, Pvalb) cell, we trained classifiers on a subset of the dataset and evaluated their prediction accuracy on held-out data using a stratified k-fold bootstrap procedure (see Methods). Based on the apparent overlap in presynaptic cell-class–specific distributions of synaptic dynamics (Figure 2C) and the presumed alignment between these classes and their characteristic dynamics,^38^ we considered three possibilities: 1) the distributions for Pvalb-pre, Sst-pre, and VIP-pre dynamics overlap so extensively that the classes cannot be distinguished; 2) all three classes share a common set of dynamics, but some dynamics may be unique to a single class while others are shared only between pairs of classes; 3) all three classes share dynamics, but these dynamics form a linear continuum from one class to the next (Figure 3A).

**Figure 3.**
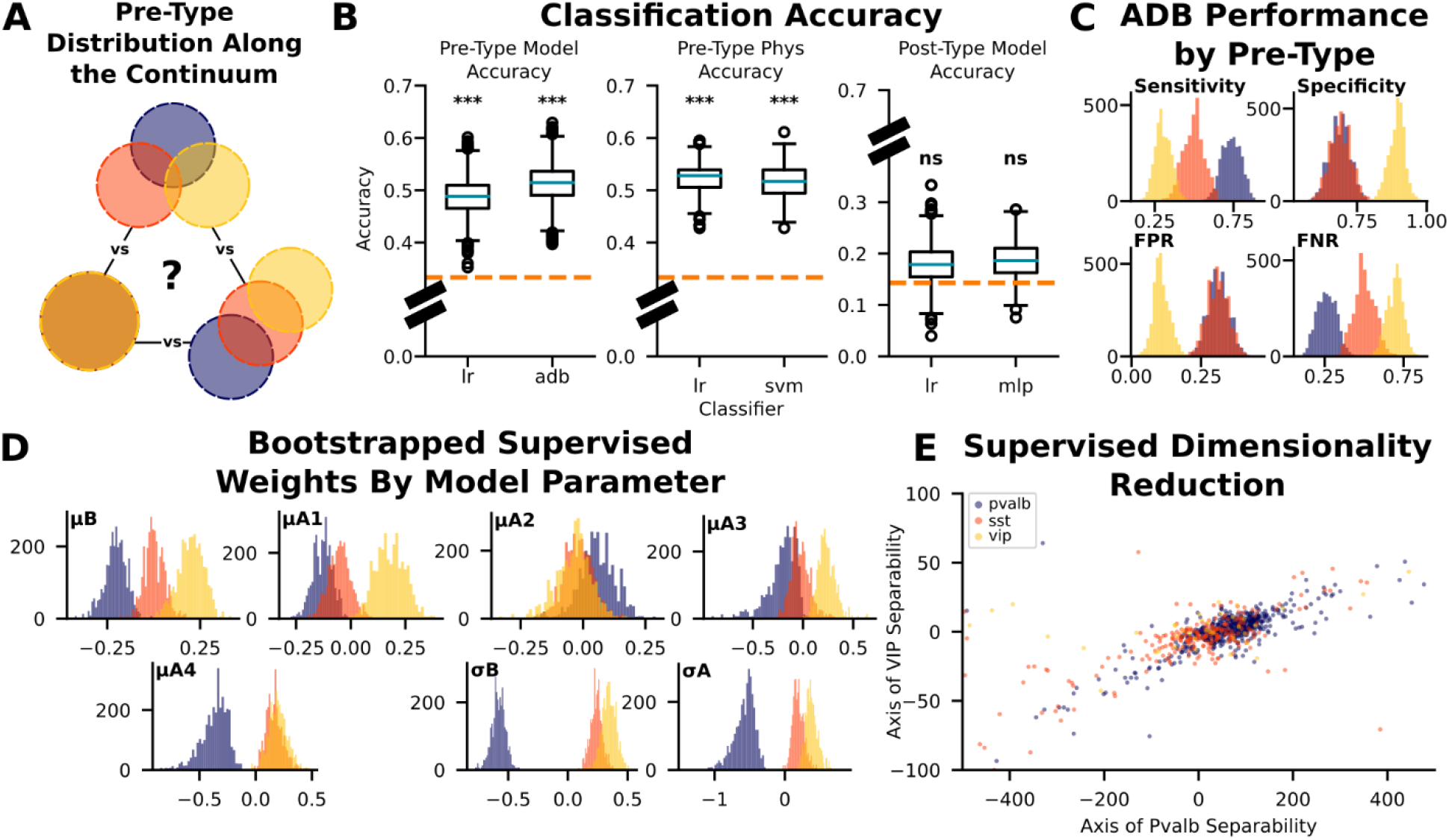
Alignment between the synaptic dynamics and the presynaptic genetic identities. **A**. Schematic of three hypothetical scenarios: 1) the distributions for Pvalb-pre, Sst-pre, and VIP-pre dynamics overlap so extensively that the classes cannot be distinguished; 2) all three classes share a common set of dynamics, but some dynamics may be unique to a single class while others are shared only between pairs of classes; 3) all three classes share dynamics, but these dynamics form a linear continuum from one class to the next. **B**. Accuracy distributions of supervised prediction of presynaptic (left, middle) or postsynaptic (right) identities based on SRP model (left, right) or physiological (middle) parameters. Best performers: “lr”: logistic regression, “adb”: Adaboost, “svm”: support vector machine, “mlp”: multilayer perceptron. **C**. Histograms of performance metrics of the Adaboost classifier (best-performing classifier using model parameters) in predicting the presynaptic identity. “FPR”: false positive rate, “FNR”: false negative rate. **D**. Histograms of the supervised weights of each model parameter used by the logistic regression to predict the presynaptic identity based on model parameters. “μ”: efficacy parameters (i.e., baseline and 4 amplitudes), “σ”: variance parameters (i.e., baseline and an amplitude). **E**. Projection of individual fits onto the space capturing the most separability between the classes, according to logistic regression (B, left).

We show that while supervised learning on both model parameters and physiological data yielded similar results, prediction accuracy was only significantly greater than baseline prediction (i.e., 1 / number of classes) when classifying the presynaptic cell type, not the postsynaptic one (presynaptic prediction using model parameters: LR accuracy=0.49±0.03, p=0.00; ADB accuracy=0.51±0.03, p=0.00; presynaptic prediction using physiological features: LR accuracy=0.52±0.03, p=0.00; SVM accuracy=0.52±0.03, p=0.00; postsynaptic prediction using model parameters: LR accuracy=0.18±0.04, p=0.19; MLP accuracy=0.19±0.03, p = 0.11) (Figure 3B). Thus, inhibitory synaptic dynamics tended to align more closely with the genetic identity of the presynaptic cell, contrasting with excitatory dynamics showing a stronger association with the identity of the postsynaptic cell.^38,40^ Also, when evaluating the best-performing classifier (Adaboost^47^), we found it to be most accurate in identifying Pvalb-pre synapses, showing the highest true positive rate among the three types. In comparison, it frequently misclassified VIP-pre synapses, most often labeling them as Sst-pre synapses (Figure 3C). The combination of these results and the significant prediction accuracies indicate an overlap between the spread of dynamics of these three genetic classes, wherein Pvalb-pre synapses may display the more consistent or distinct ones.

To determine which model features were most important for distinguishing among presynaptic subclasses, we quantified feature importance by extracting, at each bootstrap iteration, the average predictive weight assigned to each model parameter by the logistic-regression classifier.^47^ Figure 3D, corresponding to histograms of these supervised weights by model parameter for each presynaptic class, shows that the variance parameters (σ) are the ones that the logistic regression finds the most informative in distinguishing Pvalb-pre synapses from the others. Among the efficacy kernel parameters, the amplitudes of the slow components (μA3 and μA4) also contribute strongly to the classifier’s ability to distinguish Pvalb-pre synapses from the others. These results can be visualized in Figure 3E, which shows rodent inhibitory synaptic dynamics projected into a simplified space where each axis reflects the directions that best separate the different presynaptic subclasses, according to the logistic regression. To further this inquiry to physiological parameters, we repeated the same analysis using a set of physiological measures instead of model parameters (see Methods). Release probability emerged as one of the most informative features in the classification, with Pvalb-pre synapses showing the highest values, followed by Sst-pre and VIP-pre synapses (Figure S1).

Therefore, although the synaptic dynamics of inhibitory synapses align more strongly with the genetic identity of the presynaptic cell than with that of the postsynaptic cell, the class-specific distributions of these dynamics remain highly overlapping, with Pvalb-pre synapses showing the greatest separability from the other two types.

### Model parameters organize into an overlapped continuum between Pvalb-pre and VIP-pre synapses with Sst-pre synapses acting as a transitional class

To gain a clearer picture of this continuum in terms of overlapping genetic subclasses of synaptic dynamics, we explored which dynamics are found in which presynaptic cell genetic subclasses, population wide. Figure 4A illustrates, for each presynaptic subclass, the predicted responses of individual synapses to their stimulation protocols (left column) and how the predicted standard deviation of those responses evolves across pulses (right column). As reported in the literature,^38^ Pvalb-pre synapses exhibit the strongest short-term depression, the lowest response variability, and a consistent decline in response variability across successive pulses – as would be typically expected from high release probability synapses. This is reflected in their medoid efficacy kernel (i.e., the kernel corresponding to the most centrally located exemplar within the cluster) (Figure 4B, left) and variance kernel (Figure 4B, right). The general consistency of short-term behaviors and distinctiveness, especially in terms of variance profiles (Figure 4C), likely explains its greater classification accuracy. As for Sst-pre and VIP-pre synapses, both tend to express a mixture of biphasic, depressing and facilitating synapses, while displaying greater, yet still declining, variability in responses. Interestingly, the Sst-pre medoid^47^ efficacy kernel (Figure 4B) shows short-timescale facilitation and longer-timescale depression, indicative of responses that depend strongly on input frequency.

**Figure 4.**
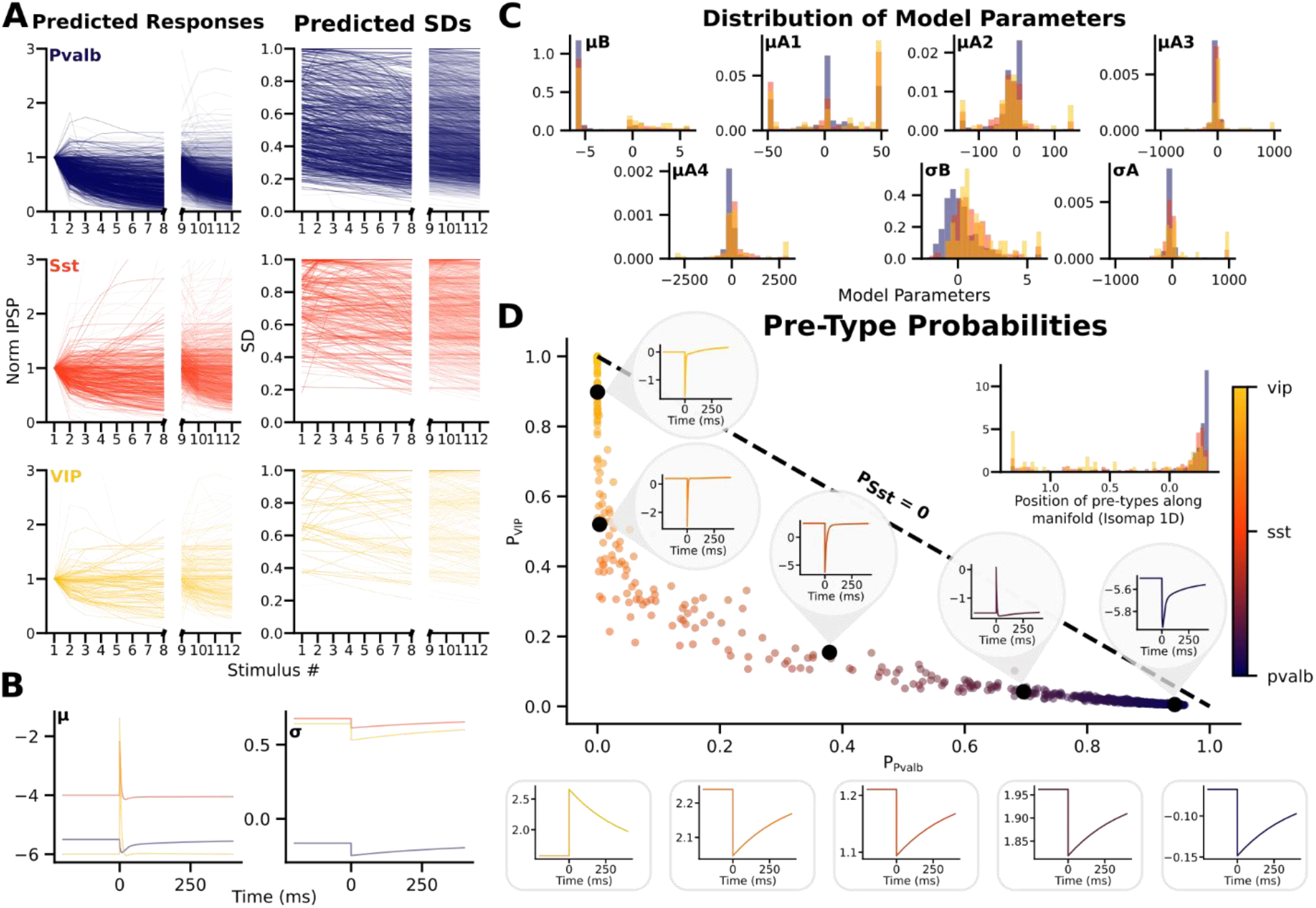
Short-term Dynamics Along the Continuum across Pvalb-pre, Sst-pre and VIP-pre synapses. **A**. All predicted responses (left) and predicted SDs (right) by presynaptic cell subclass. **B**. Medoid efficacy and variance kernels by presynaptic cell subclass. **C**. Histograms of the model parameters across presynaptic cell subclasses. “μ”: efficacy parameters (i.e., baseline and 4 amplitudes), “σ”: variance parameters (i.e., baseline and an amplitude) **D**. Naive Gaussian Bayes–based probabilistic assignment of short-term dynamics to the VIP and Pvalb classes. Within-graph efficacy kernels are medoids obtained from k-means clusters separating the probability distributions into five groups. Out-of-graph variance kernels are color-coded to correspond to their within-graph efficacy kernel. Subplot illustrates the true distribution of synapses along this continuum, wherein the two-dimensional probability outputs of the classifier were reduced to a single continuous axis using Isomap and then represented in the form of a density-based histogram.

Following this analysis, we sought to establish how dynamics change along the continuum. To this end, we trained a Gaussian Naive Bayes classifier^47^ on the rodent dataset following the same protocol as for the others described before. For each synapse, we then extracted the predicted probability that it belongs to each of the three presynaptic cell subclasses. Figure 4D plots these results, with one axis representing the predicted probability that a synapse is Pvalb-pre and the other representing the probability that it is VIP-pre. This positions Sst-pre synapses as a transitional class between Pvalb-pre and VIP-pre synapses: no dynamics are predicted to belong equally to the Pvalb-pre and VIP-pre subclasses without also showing a strong probability of being Sst-pre, and none are predicted to belong almost exclusively to the Sst-pre subclass. We then applied k-means clustering to these sets, resulting in five distinct groups each. Computing the medoid efficacy kernels of the corresponding synapses, we observed a smooth transition between VIP-pre and Pvalb-pre quasi-exclusive dynamics. In the rodent dataset, this progression spans from synapses showing a mixture of fast-timescale depression and longer-timescale facilitation (yellow), to short-timescale depressing synapses (orange to red-orange), followed by fast-timescale facilitating and longer-timescale depressing synapses (purple) to strongly depressing synapses (dark blue). In terms of response variance, we observe a continuum ranging from synapses with high and increasing variability across pulses (yellow) to those whose variability steadily decreases over the train, both in average magnitude and pulse-to-pulse fluctuations (orange to dark blue).

Although the Gaussian Naive Bayes classifier suggests a continuum running from VIP-pre to Pvalb-pre synapses, with dynamics at either extreme predicted to belong almost exclusively to those respective subclasses, the true distribution of synapses along this continuum is more nuanced. To visualize this, the two-dimensional probability outputs of the classifier were reduced to a single continuous axis using Isomap^47^ (subfigure of Figure 4D). This projection reveals how synapses from each class are truly distributed along the continuum, independent of the classifier’s predictions. Even though each class is most abundant near the end of the continuum (i.e., where the classifier predicts Pvalb-pre synapses), the enrichment is not uniform: Pvalb-pre synapses are disproportionately concentrated at this end, followed by Sst-pre and then VIP-pre synapses. In contrast, the remainder of the continuum is occupied almost exclusively by Sst-pre and VIP-pre synapses, with VIP-pre synapses becoming proportionally more common toward the opposite end (i.e., where the classifier predicts VIP-pre synapses). Overall, there exists an overlapped continuum between Pvalb-pre and VIP-pre synapses with Sst-pre synapses acting as a transitional class.

## Discussion

Using a combination of machine learning and synaptic modeling on the largest existing paired-patch clamp synaptic dynamics dataset we show that the dynamics of inhibitory connections organize into a functional continuum with distinct but overlapping modes corresponding to different genetic identities. Unlike in excitatory synapses,^40^ unsupervised machine learning did not yield well defined clusters. Instead, both our model parameters and supervised prediction weights are organized into a relatively smooth continuum between Pvalb-pre and VIP-pre synapses with Sst-pre synapses acting as a bridging, transitional class.

This distinction in the organization of excitatory and inhibitory dynamics raises the question of why these populations would differ so fundamentally; in other words, what functional roles may be attributed to excitatory subtypes compared to an inhibitory continuum. One possibility can be drawn from previous work suggesting that STP can act as a temporal reservoir, with heterogeneous synaptic dynamics providing a set of basis functions. Indeed, Barri et al. demonstrated that diverse STP dynamics at mossy fiber-granule cell connections in the cerebellum results in superior processing of sub-second time representations.^30^ In this framework, heterogeneity in synaptic dynamics allows the network to represent a richer set of temporal features. Therefore, a continuum of inhibitory STP may provide a flexible means of shaping the temporal flow of activity within a circuit. This aligns with the idea that inhibitory interneurons act as dynamic stabilizers, continuously adjusting their influence according to ongoing activity and preventing runaway excitation.^48,49^

Another line of theory suggests that STP may support the multiplexing and decoding of multiplexed information streams by enabling synapses to respond selectively to specific temporal motifs. In this view, STP functions as a set of dynamic filters, enabling downstream neurons to extract distinct components from a shared input.^28,29^ Related experimental work has shown that synapse-specific STP profiles enable temporally sensitive integration of convergent multisensory inputs, with the dynamics of each input pathway encoding contextual information about its source and timing.^30^ Excitatory subtypes may be particularly well suited to this role: distinct excitatory classes could specialize for different input statistics, with their characteristic STP profiles tuned to particular temporal patterns.

Under this interpretation, functional subtypes of excitatory STP could reflect a specialization for motif-specific processing. A complimentary inhibitory continuum could reflect a graded, compensatory mechanism for regulating and shaping motif-specific computations while contextualizing them within a precise representation of time. Determining whether this division of labor truly reflects a fundamental organizational principle of cortical computation, and to what extent each form of organization reflects intrinsic synaptic properties versus learned features emerging from network-level activity, remain important questions for future experimental and theoretical work.

### Limitations

While fitting the SRP model to characterize short-term dynamics provides a precise and compact representation of the underlying patterns, the extent to which the model captures the biological responses remains a key limitation of this study. We selected the exponential decays’ time constants for both the efficacy and the variance kernels in a way that allows the model to capture a wide variety of dynamics; however, in some especially difficult cases, the model could fail to do so, resulting in sets of parameters settling on boundaries of their space. This artificially creates an area susceptible to be identified as a cluster. Likewise, since these failed fits were more common in the VIP class, it could mislead the classifiers into learning patterns that are artifacts of poor model fitting rather than meaningful biological distinctions. Therefore, we opted to remove these fits, meaning that some potentially meaningful but difficult dynamics to characterize were left out. We were also constrained to using mono-exponential decay kernels for the variance, because the model was unable to capture how variance profiles differed across stimulation protocols at individual synapses. This limitation reduced our ability to represent more complex forms of heteroskedasticity.

Regarding the continuum hypothesis testing, our results should be interpreted as supporting a continuum hypothesis rather than proving it. Indeed, the lack of significant differences between the SC distributions obtained from bootstrapped subsampling before clustering on the biological data and those from the synthetic data indicates that our clusters are not of substantially higher quality than those that could arise from clustering a multivariate Gaussian with a comparable variance structure; in other words, the inhibitory clusters obtained using OPTICS are not meaningful, supporting the notion of a continuum without proving it. That said, applying a similar method onto excitatory dynamics yielded opposite results, highlighting potential differences between the two groups; excitatory dynamics may be more inclined to form subtypes whereas inhibitory ones may occupy a more continuous region of parameter space. Still, of note are the procedural differences between the two groups: we used fewer excitatory (86 synapses) than inhibitory (940 synapses) synapses and we constructed the efficacy kernel for the excitatory datasets using 4 parameters compared to 5 for the inhibitory ones. These differences can impact the SCs, partly explaining the greater SC associated with the excitatory dynamics. However, the contrasting results from the continuum hypothesis testing help mitigate these concerns, and we therefore believe that these limitations do not substantially weaken the overall conclusion.

## Methods

### Spike response plasticity model

We used the Spike Response Plasticity model,^44,45^ a phenomenological model that forgoes some biophysical interpretability to extend its flexibility, thus aligning well with the goal of this project. Precisely, this model constructs a kernel through a linear combination of basis functions (here, exponential decay functions), each governed by model parameters. For an exponential decay function, these parameters include a time constant (not trainable) and an amplitude (trainable). This kernel is then convolved with a stimulus train (i.e., the 12 pulses), to which is added a baseline parameter (trainable). To account for intrinsic synaptic nonlinearities (e.g., the saturation of release probability), the result is passed through a sigmoidal readout function. The amplitudes were normalized to the first pulse, such that this value is divided by the nonlinear readout of the baseline. The resulting trace could be interpreted as the time-dependent synaptic efficacy, where a succession of spikes could lead to gradually increasing/decreasing (i.e., with (+)/(-) amplitudes) efficacies at spike times, while between spikes, efficacies would decay. A schematic of this process is shown in Figures 1A and 1B.

In this model, there are two different kernels: an efficacy kernel (constructed as a sum of 4 exponential decay with time constants: 5 ms, 15 ms, 200 ms, and 4000 ms) and a variance kernel (constructed as a mono-exponential decay with a time constant of 400 ms). Therefore, a total of seven parameters (i.e., 5 amplitudes and two baseline parameters) are fitted to recorded responses for each synapse.

### Data Processing

We restricted our analysis to current-clamp recordings performed in 1.3 mM extracellular calcium and 1 mM magnesium, as these were the conditions available that matched the most closely real physiological ones. We also restricted our analysis to pairs in which the presynaptic cell was labeled as a GABAergic interneuron (Pvalb, Sst, or VIP). Quality control excluded runs containing spontaneous action potentials or clear experimental artifacts, and we retained only traces that passed the dataset’s built-in quality check (qc_pass).^38^ This quality control required, among other criteria, low baseline noise (<5 mV), no additional presynaptic pulses within ±8 ms of the stimulus, an inhibitory baseline membrane potential between −60 mV and −50 mV, a postsynaptic recording standard deviation below 1.5 mV, and a maximal response amplitude below 10 mV.

We pooled runs with small variations in delay duration (125–128 ms and 250–253 ms) and padded missing responses with NaNs. Postsynaptic potential amplitudes were computed by subtracting the mean membrane voltage in the 2 ms window preceding the spike from the mean voltage in the 2–10 ms window following it. Responses with amplitudes below −0.005 V were excluded, and values above −10^−9^ V were clipped to −10^−9^ V to remove extreme outliers. Finally, PSP amplitudes were normalized by dividing all responses from a given synapse by the average of that synapse’s first recorded responses across protocols.

### Model Fitting

Assuming responses at each pulse over multiple trials followed a Gaussian distribution, we used the mean squared error (MSE) as the loss function. To minimize the loss, we used simplicial homology global optimization as the algorithm.^50^ Furthermore, to increase the probability of finding the global minimum of our loss, we repeated the optimization over multiple intervals that collectively spanned the full range of the baseline parameter space (i.e., from -6 to 6 divided in twelve intervals of unit length). The overall fitting procedure went as follows: first, the five parameters associated with the efficacy kernel were fitted by minimizing the MSE between the observed responses and the model’s predictions. Then the last two parameters corresponding to the variance kernel were fitted, but between the observed standard deviation of responses at each pulse and the predicted standard deviation of responses.

### Unsupervised Learning

To determine whether subtypes of synaptic dynamics could be extracted from the dataset, we use the density-based clustering algorithm OPTICS. Since the dataset is heavily class-imbalanced, we used a bootstrap subsampling before clustering methodology. This involved sampling without replacement from the dataset to retrieve an equal number of synapses associated with each presynaptic subclass (i.e., for a total of 204 rodent synapses used for the clustering at each iteration). For this bootstrap clustering, only the efficacy kernel parameters were used to characterize synapses (i.e., a baseline parameter and four amplitudes) as they are the ones conveying information on the short-term behavior of the mean efficacy.

First, we used Principal Component Analysis (PCA) at each iteration of the bootstrap to transform the original feature space into a new space where, by definition, variability is maximized along each principal component. Then, we clustered the subset of the dataset while trying a range of minimum cluster sizes, a hyperparameter OPTICS uses to control the granularity of cluster detection by defining the smallest number of points required to form a dense region. Clustering quality was assessed by computing a Silhouette Coefficient (i.e., the average, over all data points, of the quantity (b_i_-a_i_)/max(a_i_, b_i_), where a_i_ is the mean intra-cluster distance, and b_i_ is the mean nearest-cluster distance). The optimal clustering for a certain subset of the dataset was selected as the one yielding the highest Silhouette Coefficient. Also, to determine whether subtypes of synaptic dynamics can be identified within presynaptic subclasses separately, clustering was performed on each subclass separately. The same steps were performed independently on the layer-specific subsets of the total dataset.

### Continuum Hypothesis Testing

Presumably, because OPTICS identifies clusters where there is greater density of data points, it should always be predicting the presence of at least two groupings (e.g., one “real” cluster, and one “noise” cluster), regardless of whether we expect the dataset to form groupings or not. Therefore, to test the significance of potential observed clusters, we tested the biological data against a continuum hypothesis (i.e., the hypothesis that there aren’t significant groupings in the dataset). To do this, we generated 204 synthetic synapses by sampling from a multivariate Gaussian whose variance structure matches the variance structure of the observations. Specifically, we defined this distribution such that its primary axis of variance corresponds to the one captured by the first principal component (PC1) of the biological data, while the remaining variance was distributed evenly between the other four dimensions. This created a natural axis onto which artificial clusters can form. These samplings were performed 3000 times such that presynaptic cell types were equally represented in biological samples and that the size of the synthetic samples corresponded to the size of the biological ones. During each iteration, SCs were computed from OPTICS applied on both biological and synthetic data (separately) for a series of minimum cluster sizes yielding exactly two clusters. Thus, each iteration yielded two SCs corresponding to the highest quality clustering of the biological and synthetic data, respectively. P-values were computed as the proportion of iterations in which the biological silhouette coefficients exceeded the corresponding synthetic silhouette coefficients.

### Supervised Learning

To assess whether features of synaptic dynamics—captured either by the SRP model parameters or by a set of physiological measurements—can predict the transgenic class of the presynaptic (i.e., Pvalb, Sst, VIP) or postsynaptic (i.e., Nr5a1, Tlx3, Sst, Ntsr1, VIP, Sim1, Pvalb) cell, we trained classifiers on a subset of the dataset and evaluated their prediction accuracies on held-out data. The SRP model parameters consisted of the five parameters associated with the efficacy kernel (i.e., a baseline and four amplitudes) and the two parameters associated with the variance kernel (i.e., a baseline and one amplitude). The physiological features included the mean summed value of the first eight stimuli, the mean ratio of the fifth response over the first response, the second over the first (for all runs with a 50 Hz induction frequency), the release probability, and the mean value of the last four responses in a train (these responses divided by the mean of the first four for all protocols). We considered 6 choices of classifiers: gradient boosting, AdaBoost, logistic regression, random forest, multilayer perceptron, and support vector machines.^46^

As the distribution of synapses across cell classes is imbalanced, risking classifier bias toward larger classes, we use a stratified k-fold bootstrap method with per-fold class balance. This method divides the initial 940 synapses into *k* (here, *k* = 5) sets, maintaining class proportions as in the original dataset. By rotating through these sets, it then selects one of these subsets to be the testing set while the remaining sets serves as the training set. At each iteration through this cycle, a different test set was selected, thus allowing each k-fold to be a part of the training set *k*-1 times and be the testing test one time. A total of 3171 bootstrap iterations were performed. To ensure class balance in the training set, we used oversampling (i.e., sampling with replacement) from minority classes (i.e., Sst and VIP when predicting the presynaptic cell class) while class balance in the testing dataset was achieved by sampling without replacement from the majority classes (i.e., Pvalb and Sst when predicting the presynaptic cell class). The prediction accuracy at each iteration (i.e., one cycle through the five subsets) corresponds to the average fraction of correct predictions over this cycle, for a total of 3171 iterations.

We selected medoid kernels using the KMedoids algorithm^47^ (with *k* = 1), defining the medoid within each synapse group (e.g., synapses sharing a presynaptic cell type) as the kernel whose total pairwise dissimilarity to all other kernels in the group was minimal. SRP model parameters were first normalized to the range 0–1 to ensure that kernel dissimilarities reflected genuine structural variation rather than differences in parameter scale.

## Acknowledgments

This work was supported by the Canadian Institutes of Health Research (CIHR) Discovery Grant RN38364 (to R.N., providing salary to J.P. and J.B.), the Brain-Heart Interconnectome Training Award from the Canada First Research Excellence Fund (CFREF) (to R.N., providing salary to J.P.) (CFREF-2022-00007), the Natural Sciences and Engineering Research Council of Canada (NSERC), Canada Graduate Research Scholarship – Master’s program (to J.P., providing salary to J.P.), Postgraduate Scholarships-Doctoral (to J.B., providing salary to J.B.), and a Canada Research Chair to K.T. The funders had no role in study design, data collection and analysis, decision to publish, or preparation of the manuscript. We also acknowledge the Creative Commons-licensed synapse image (https://doi.org/10.7875/togopic.2020.200, used with modification), Multivariate normal distribution image (CC0 1.0), function, histogram, mouse, user, dataset and dice images (‘‘image: Flaticon.com’’).

---

**Figure S1.**
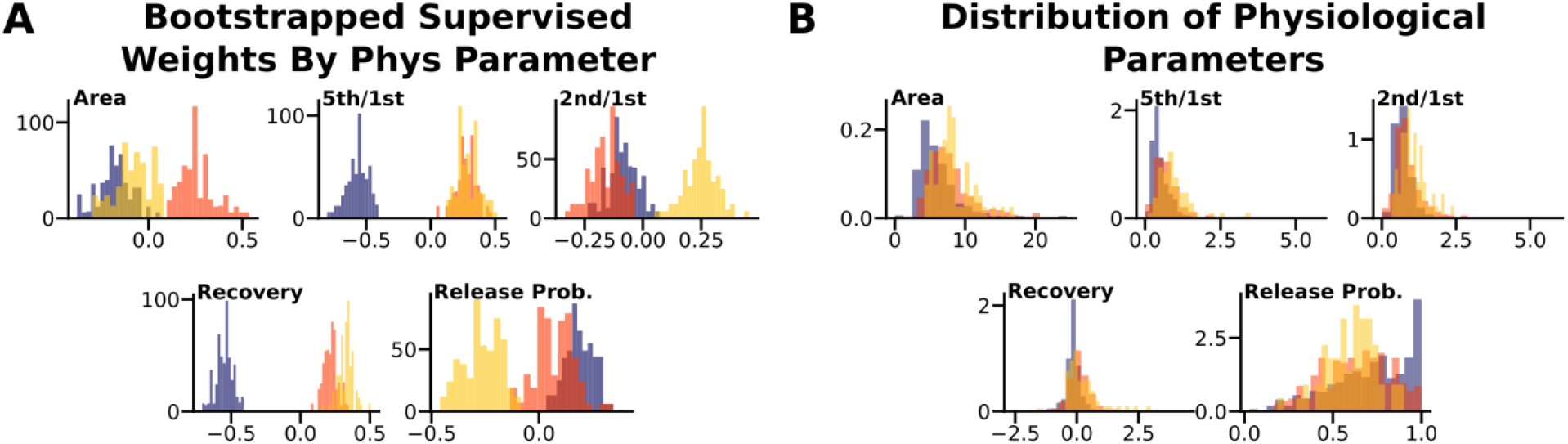
Distributions of Logistic-Regression Weights and Underlying Physiology Features Across Presynaptic Subclasses. **A**. Histograms of the supervised weights of each physiology parameter used by the logistic regression to predict the presynaptic identity. **B**. Histograms of the physiology parameters across presynaptic cell subclasses. “Area”: the mean of the summed values of the first eight responses across all trains, “5^th^/1^st^”: the mean ratio of the 5th response to the 1st response across all trains, “2^nd^/1^st^”: the mean ratio of the 2nd response to the 1st response across all trains, “Recovery”: the mean of the last four responses in each train divided by the mean of the first four responses, averaged across all protocols.

